# Elevated oxidative phosphorylation is critical for immune cell activation by polyethylene wear particles

**DOI:** 10.1101/2022.10.19.512774

**Authors:** Chima V. Maduka, Maxwell M. Kuhnert, Oluwatosin M. Habeeb, Anthony Tundo, Ashley V. Makela, Stuart B. Goodman, Christopher H. Contag

**Affiliations:** Comparative Medicine & Integrative Biology, Michigan State University, East Lansing, MI 48824, USA; Department of Biomedical Engineering, Michigan State University, East Lansing, MI 48824, USA; Institute for Quantitative Health Science & Engineering, Michigan State University, East Lansing, MI 48824, USA; Department of Orthopedic Surgery, Stanford University, CA 94063, USA; Department of Bioengineering, Stanford University, CA 94305, USA; Department of Microbiology & Molecular Genetics, Michigan State University, East Lansing, MI 48864, USA

**Keywords:** Polyethylene wear particles, macrophage, mitochondrial oxidative phosphorylation, total joint replacement

## Abstract

Chronic inflammation is a major concern after total joint replacements (TJRs), as it is associated with bone loss, limited bone-implant integration (osseointegration), implant loosening and failure. Inflammation around implants could be directed away from adverse outcomes and toward enhanced osseointegration and improved surgical outcome. Activated macrophages exposed to polyethylene particles play a dominant inflammatory role, and exhibit elevated mitochondrial oxidative phosphorylation (OXPHOS) whose role is unclear. By probing the contribution of the electron transport chain (ETC), we show that increased oxygen consumption does not contribute to bioenergetic (ATP) levels in fibroblasts and primary bone marrow-derived macrophages activated by polyethylene particles. Rather, it generates reactive oxygen species (ROS) at complex I by increasing mitochondrial membrane potential in macrophages. Inhibition of OXPHOS in a dosedependent manner without affecting glycolysis was accomplished by targeting complex I of the ETC using either rotenone or metformin. Metformin decreased mitochondrial ROS and, subsequently, expression of proinflammatory cytokines, including IL-1β, IL-6 and MCP-1 but not TNF-a in macrophages. These results highlight the contribution of mitochondrial bioenergetics to activation of immune cells by polyethylene wear particles, offering new opportunities to modulate macrophage states toward desired clinical outcomes.

## 1. Introduction

Degenerative osteoarthritis is commonly treated by primary total joint replacements (TJRs). Bearing surfaces of articulations are often lined by polyethylene to prevent metal-on-metal articulation which is associated with metal ion toxicities and aseptic lymphocyte-dominated vasculitis-associated lesions^1^. However, wear particles of polyethylene generated by relative motion of artificially reconstructed joints after TJRs activate macrophages, resulting in chronic inflammation and bone loss^2,3^. This limits bone-implant integration (osseointegration) necessary for implant longevity^4,5^. Implant longevity is crucial because revision surgeries are costlier, more difficult and associated with higher complication rates than primary TJAs. Crosslinking polyethylene increases its resistance to wear, significantly reducing, but not eliminating, wear particle generation^6^. Wear particles of ultrahigh molecular weight polyethylene (UHMWPE) or highly crosslinked polyethylene (XLPE) are clinically relevant and result in inflammation^6,7^. Advances in understanding the role of macrophage polarization in macrophage-mesenchymal stem cell crosstalk show that inflammation induced by wear particles represents an opportunity to re-direct inflammation toward desired surgical outcomes, such as enhanced osseointegration and implant longevity^8–13^.

The mitochondrion is the center of energy (adenosine triphosphate; ATP) production in the cell, meeting bioenergetic demands by oxygen consumption (oxidative phosphorylation; OXPHOS) under resting conditions. As part of OXPHOS, electron transfer through the mitochondrial electron transport chain (ETC) generates ATP^14^. The ETC consists of a series of protein complexes, including complex I, II, III, IV and V with diverse roles. Beyond meeting energy demands, the ETC plays key regulatory and effector roles in inflammation^15–18^. In neutrophils and T-cells, increased oxygen consumption results in superoxide formation. Although oxygen consumption is reduced in macrophages exposed to bacterial lipopolysaccharide (LPS) as part of the switch from OXPHOS to glycolysis for ATP production^19,20^, residual oxygen consumption is directed to superoxide formation^19^, eventually producing reactive oxygen species (ROS)^21^ as part of inflammation.

The immunometabolic changes in macrophages exposed to UHMWPE wear particles includes elevated OXPHOS^22^, but its precise role is unclear. Herein, we set out to probe the role of increased oxygen consumption in macrophages activated by polyethylene particles. Elucidating the role of mitochondrial bioenergetics in immune cells activated by polyethylene particles could offer new insights toward controlling inflammation around implants. Clinically, this could translate to enhanced osseointegration, improved surgical outcome and implant longevity.

## 2. Materials and methods

### 2.1. Cells

Primary bone-marrow derived macrophages (BMDMs) derived from C57BL/6J mice (Jackson Laboratories) of 3-4 months^19,23^ as well as wild-type mouse embryonic fibroblast (MEFs) cell line (NIH 3T3 cell line; ATCC) which tested negative for Mycoplasma were used. Cells were seeded in complete medium comprising DMEM medium, 10% heat-inactivated Fetal Bovine Serum and 100 U/mL penicillin-streptomycin; all reagents were sourced from ThermoFisher Scientific. For the different assays, 50,000 BMDMs or 20,000 MEFs were seeded for 3 days in a 96-well tissue culture plate in 200 μL of complete smedium.

### 2.2. Materials

The generation, characteristics and concentration (1.25 mg/ mL) of endotoxin-free UHMWPE particles used in this study have been previously described^7^. Additionally, 100ng/ mL of lipopolysaccharide (LPS) from *Escherichia coli* O111:B4 (MilliporeSigma) was used. Reagents including rotenone, metformin, oligomycin and antimycin A were sourced from MilliporeSigma and used at concentrations shown in each figure for probing mitochondrial functions. These reagents were reconstituted in complete medium and added at the time of seeding cells for experiments.

### 2.3. Bioenergetic (ATP) measurement

Bioenergetics was measured using ATP/ADP kits (Sigma-Aldrich) according to manufacturer’s instructions. Luminescence was measured using the SpectraMax M3 Spectrophotometer (Molecular Devices) by SoftMax Pro (Version 7.0.2, Molecular Devices) at integration time of 1,000 ms.

### 2.4. Seahorse assay

Basal measurements of oxygen consumption rate (OCR), extracellular acidification rate (ECAR) and lactate-linked proton efflux rate (PER) were acquired using the Seahorse XFe-96 Extracellular Flux Analyzer (Agilent Technologies)^19,24,25^. The Seahorse XF DMEM medium (pH 7.4) was used after supplementation with 25 mM D-glucose and 4 mM Glutamine; seeded cells were washed off and incubated in a non-CO2 incubator an hour prior to the assay, according to manufacturer’s instructions for the Seahorse ATP rate assay. Afterwards, wave software (Version 2.6.1) was used to export Seahorse data directly as means ± standard deviation (SD).

### 2.5. Crystal Violet assay

The crystal violet assay was used to assess for cell viability^26^. Absorbance (optical density) measurements at 570 nm were acquired using the SpectraMax M3 Spectrophotometer (Molecular Devices) and SoftMax Pro software (Version 7.0.2, Molecular Devices).

### 2.6. Milliplex assay

Cytokine and chemokine expression in cell culture supernatants were evaluated using a MILLIPLEX MAP mouse magnetic bead multiplex kit (MilliporeSigma)^27^. Assays were performed for IL-6, MCP-1, TNF-a, IL-1β, IL-4, IL-10, IFN-λ and 1L-13. Data was acquired using a Luminex 200 instrument (Luminex Corporation) and xPONENT software (Version 3.1, Luminex Corporation).

### 2.7. Flow cytometry

Macrophages were detached from 96-well plates after incubation using 1X PBS with 4mM EDTA (Teknova) at 37 °C for 10 minutes, followed by gentle pipetting. Cells were assessed for mitochondrial mass (MitoTracker Green FM, MTG; ThermoFisher Scientific), mitochondrial membrane potential (tetramethylrhodamine methyl ester, TMRM; ThermoFisher Scientific) and mitochondrial superoxide (MitoSOX Red, MSOX; ThermoFisher Scientific). Unstained cells from each condition, singlestained TMRM, MTG and MSOX cells were used for controls. A mix of 1X PBS (ThermoFisher Scientific) containing Live/Dead Blue (1:1000) and MTG (50 nM)/TMRM (20 nM) or MTG (50 nM)/MSOX (2.5 uM) were added to samples containing 650,000 cells in 100 μL, followed by incubation at 37 °C and 5% CO2 in the dark for 30 min. Cells were collected by centrifugation followed by two washes using flow buffer. Flow buffer comprised 0.5% bovine serum albumin (MilliporeSigma) made in 1X PBS. Cells were fixed using 4% paraformaldehyde for 10 minutes at room temperature in the dark. Cells were collected by centrifugation followed by resuspension in 100 μL flow buffer for analysis using the Cytek Aurora flow cytometer. Cells were identified and singlets gated using FSC/SSC. MTG+ cells were gated from live cells and MSOX/TMRM were identified from the MTG+ population. Data were analyzed using FCS Express software (De Novo Software; version 7.12.0005).

### 2.8. Statistics and reproducibility

Data were presented as mean with standard deviation (SD) and analysed using statistical software (GraphPad Prism, version 9.3.1). Although exact p-values were presented, significance level was set at p < 0.05. Specific details of statistical tests and sample sizes (biological replicates) are provided in figure legends. Exported data (mean, SD) from Wave in Seahorse experiments had the underlying assumption of normality and similar variance, and thus were tested using corresponding parametric tests as indicated in figure legends.

## 3. Results

We have previously uncovered a role for glycolytic reprogramming in the inflammatory response to UHMWPE wear particles. Specific inhibition of different steps in glycolysis prevented the expression of proinflammatory cytokines while stimulating antiinflammatory cytokines^22^. However, in addition to increased glycolysis, elevated mitochondrial oxygen consumption was observed whose role was unclear.

To elucidate the role of mitochondrial OXPHOS, the function of putative sites of oxygen consumption^28^, including complex I and III of the ETC were probed, and compared to complex V where ATP is synthesized. Inhibition of complex I was accomplished using rotenone^28^; for clinical translatability, metformin which has similar pharmacologic action was also used^29,30^. Additionally, complex III and V were inhibited by antimycin A and oligomycin, respectively.

Macrophages and fibroblasts are dominant cellular actors in polyethylene particle-induced chronic inflammation and associated pathologies^3^. Compared to respective controls, exposure of primary bone marrow-derived macrophages or mouse embryonic fibroblasts to polyethylene particles decreased bioenergetics in lysed cells (Fig. 1a-b)^22^. Importantly, specific inhibition of complex I by rotenone or metformin did not further decrease ATP levels (Fig. 1a-b) in immune cells exposed to polyethylene particles. Similarly, complex III and V did not appear to contribute to ATP production in exposed immune cells (Fig. 1a-b).

**Figure 1.**
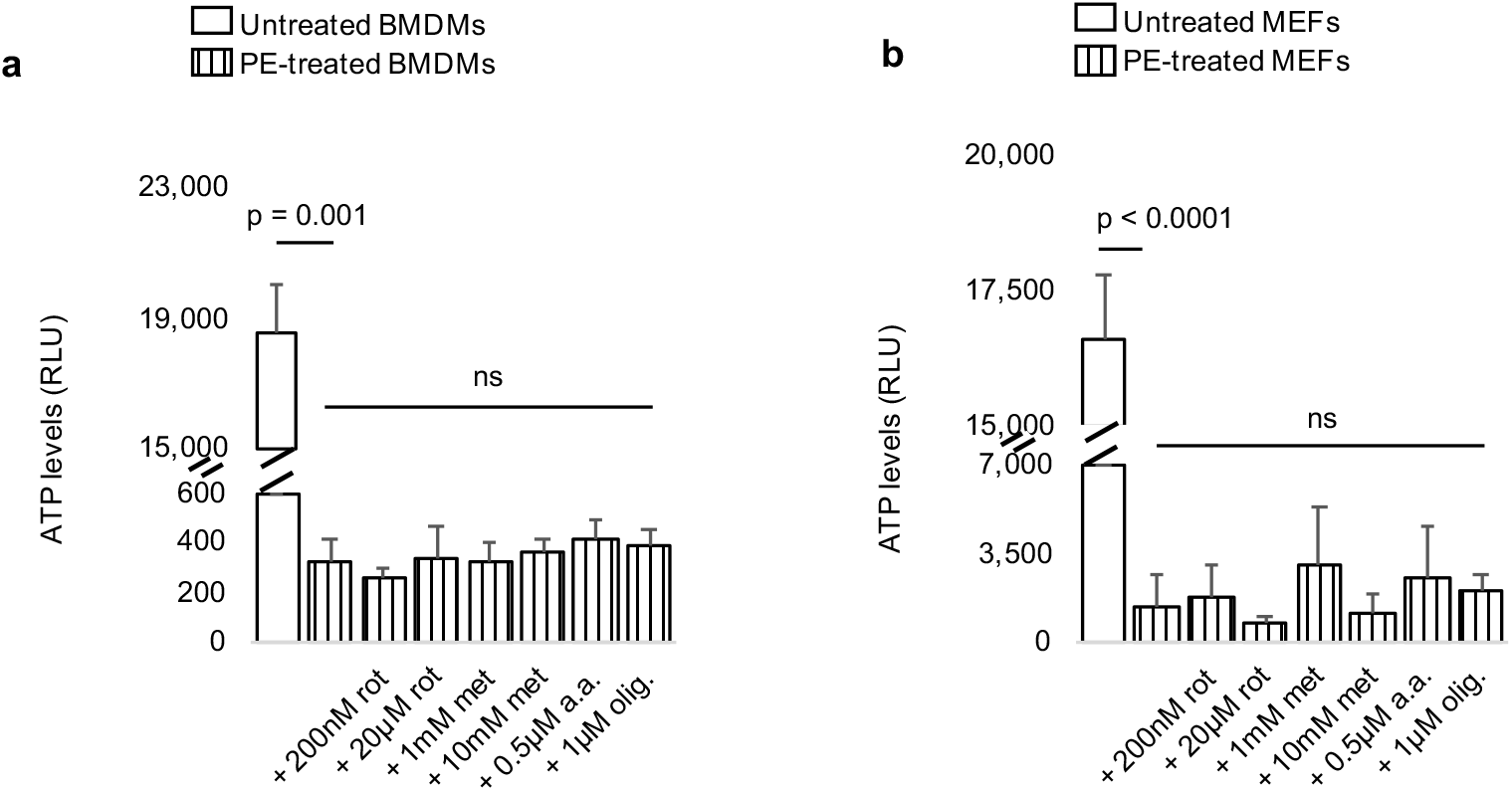
Decreased bioenergetic (ATP) levels in immune cells exposed to ultrahigh molecular weight polyethylene (PE) particles are not affected by pharmacologic inhibition of mitochondrial respiration. **a-b,** Primary bone marrow-derived macrophages (BMDMs; **a**) or mouse embryonic fibroblasts (MEFs; **b**) have decreased ATP levels after exposure to ultrahigh molecular weight polyethylene (PE) particles; decreased ATP levels are unaffected following inhibition of the electron transport chain by rotenone (rot), metformin (met), antimycin A (a.a) or oligomycin (olig.). Not significant (ns), mean (SD), n = 4 (Fig. 1a), n = 5 (Fig. 1b), Brown-Forsythe and Welch ANOVA followed by Dunnett’s T3 multiple comparisons test or one-way ANOVA followed by Tukey’s post-hoc test.

Exposure of macrophages to polyethylene particles resulted in simultaneous elevation of extracellular acidification rate (ECAR), lactate-linked proton efflux rate (PER) and oxygen consumption rate (OCR) (Fig. 2a-c)^22^.

**Figure 2.**
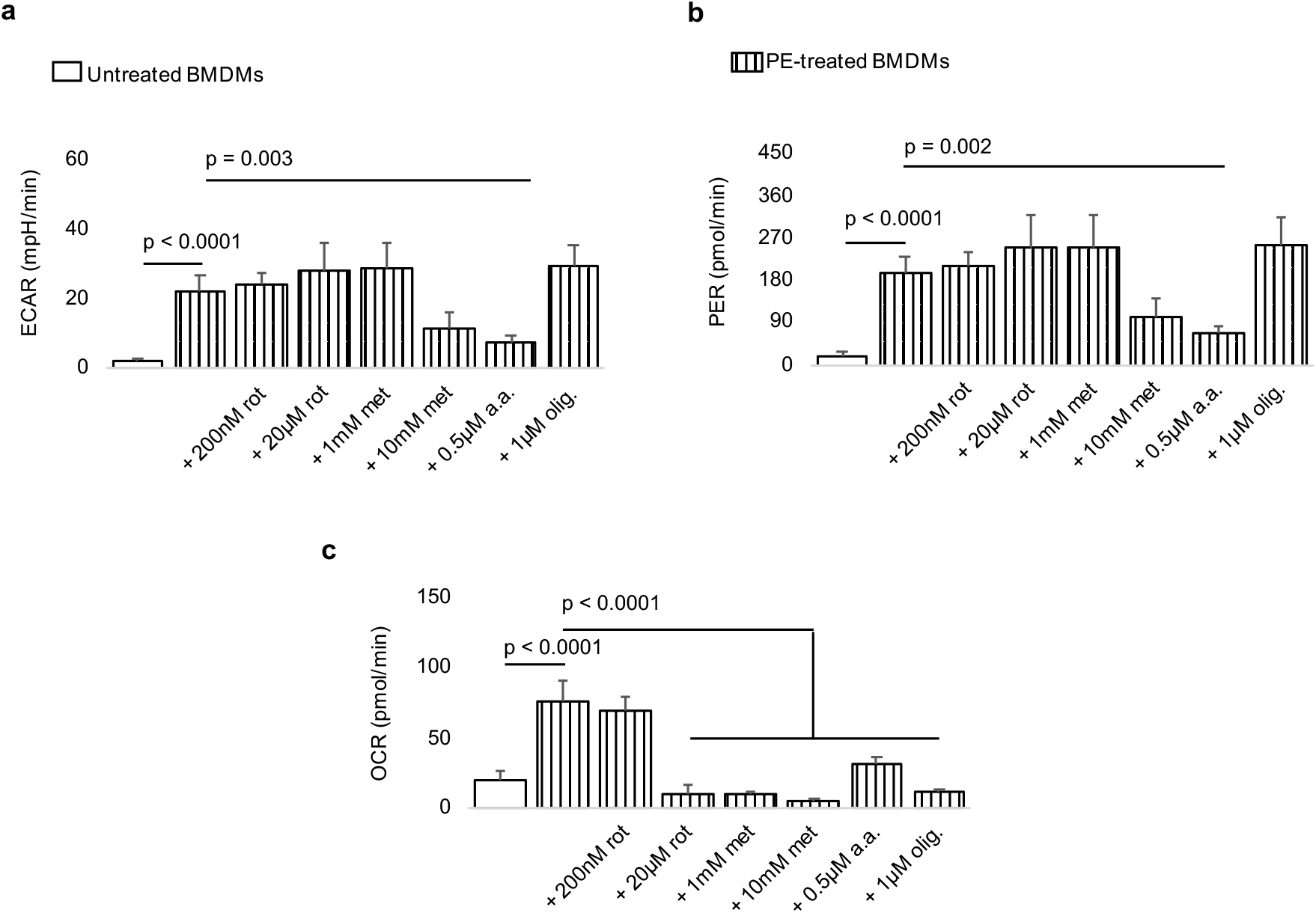
Exposure to ultrahigh molecular weight polyethylene (PE) particles increases extracellular acidification rate (ECAR), proton efflux rate (PER) and oxygen consumption rate (OCR) in macrophages; inhibitors of mitochondrial respiration reduce OCR. **a-c**, Compared to untreated primary bone marrow-derived macrophages (BMDMs), exposure to PE particles increase ECAR, PER and OCR. Inhibition of mitochondrial respiration using rotenone (rot), metformin (met), antimycin A (a.a.) or oligomycin (olig.) decreases elevated OCR but not ECAR or PER; with antimycin A, there is accompanied decrease in ECAR and PER. Mean (SD), n = 5, one-way ANOVA followed by Tukeys post-hoc test.

Whereas ECAR is indicative of glycolytic flux, OCR is an index of mitochondrial OXPHOS, and lactate-linked PER is a surrogate of monocarboxylate transporter function^31,32^, all of which are key to inflammatory activation. Compared to groups exposed to only polyethylene particles, we observed that specific inhibition of complex I and V of the ETC did not affect ECAR (Fig. 2a) or PER (Fig. 2b); however, they reduced OCR (Fig. 2c). With complex I, the reduction in OCR (Fig. 2c) was dose-dependent. On the other hand, inhibition of complex III using antimycin A reduced ECAR, PER and OCR (Fig. 2a-c) compared to groups exposed only to polyethylene particles. Importantly, pharmacological inhibition of the ETC was not the result of reduced cell numbers, excluding toxicity (Fig. 3a-b).

**Figure 3.**
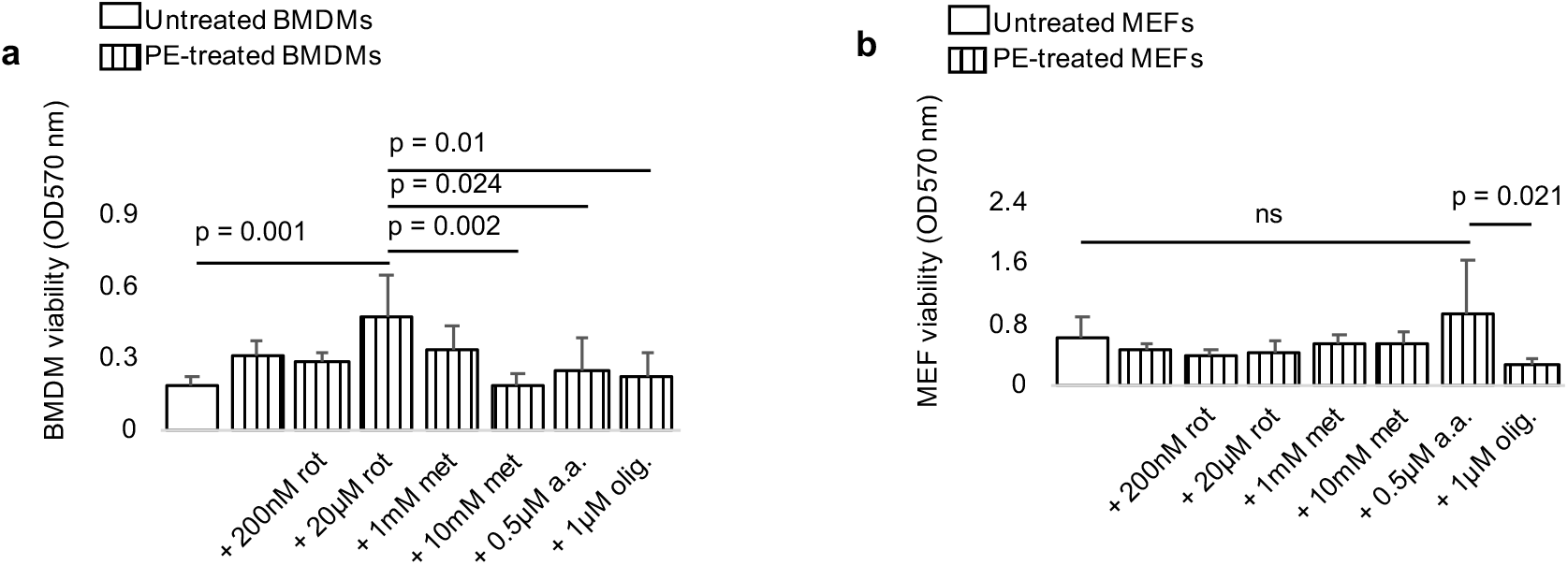
Inhibition of mitochondrial respiration does not reduce cell viability. **a-b,** In comparison to untreated cells, primary bone marrow-derived macrophages (BMDMs; **a**) or mouse embryonic fibroblasts (MEFs; **b**) have similar cell numbers after exposure to ultrahigh molecular weight polyethylene (PE) particles; cell viability is not reduced following inhibition of the electron transport chain by rotenone (rot), metformin (met), antimycin A (a.a.) or oligomycin (olig.). Not significant (ns), mean (SD), n = 5 (Fig. 1a), n = 4-5 (Fig. 1b), one-way ANOVA followed by Tukey’s post-hoc test.

To test whether increased OCR at complex I (Fig. 2c) fueled ROS production in the mitochondrion, macrophages were stained with mitoSOX Red, with a bacterial lipopolysaccharide (LPS)-treated group included for comparison. Exposure to LPS or PE particles increased mitochondrial ROS relative to untreated macrophages (Fig. 4a-c). Addition of metformin decreased mitochondrial ROS compared to macrophages exposed to only PE particles (Fig. 4d). Mitochondrial membrane potential is critical for reverse electron transport (RET) at complex I, and subsequent generation of mitochondrial ROS^19^. Tetramethylrhodamine methyl ester (TMRM) staining demonstrated that LPS or PE particles increased mitochondrial membrane potential relative to untreated macrophages (Fig. 4e-g), with metformin decreasing elevated mitochondrial membrane potential (Fig. 4h). Quantified median fluorescence intensities (MFI) corroborated these findings (Fig. 4i-j).

**Figure 4.**
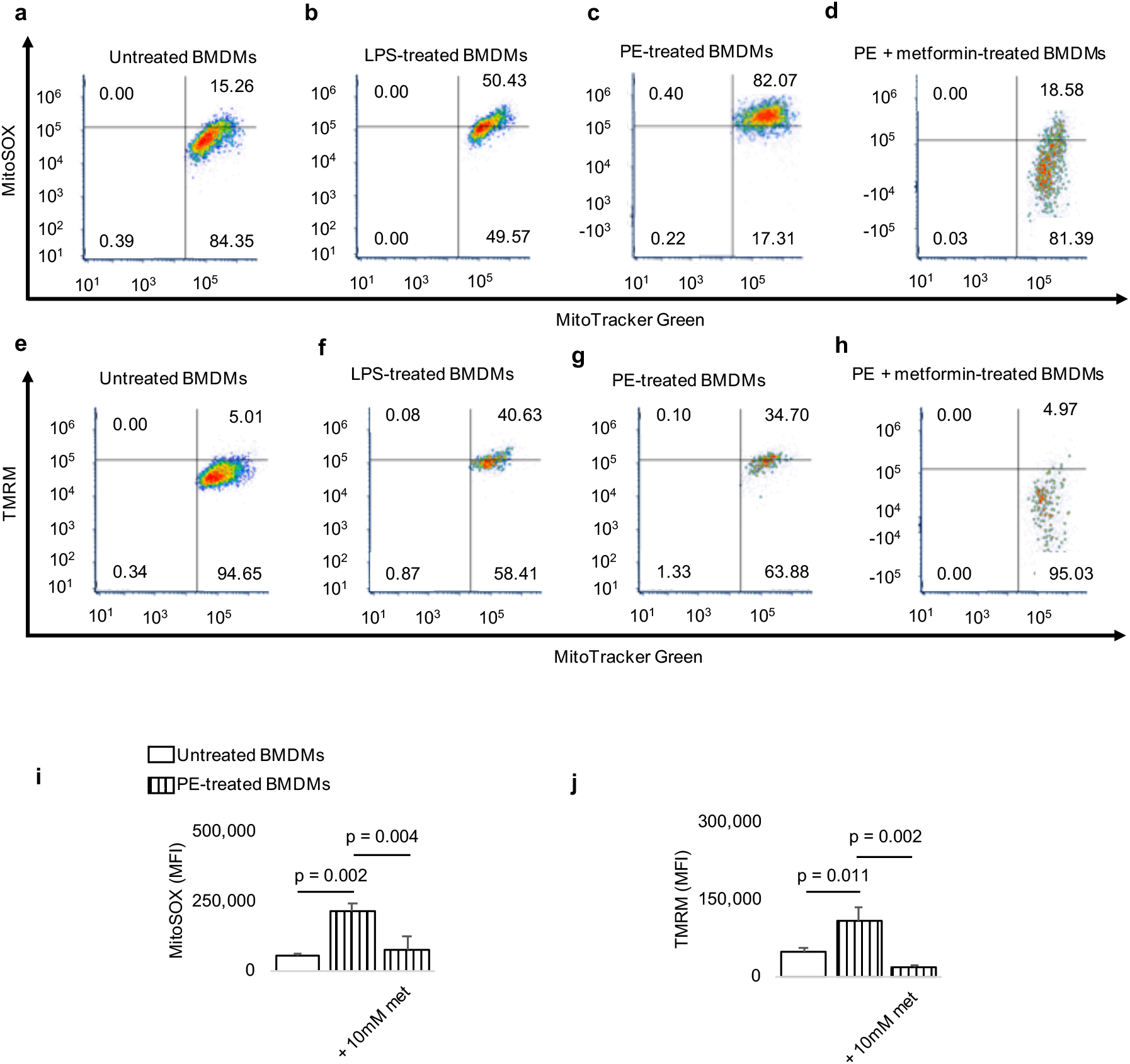
Exposure of macrophages to ultrahigh molecular weight polyethylene (PE) particles elevates mitochondrial membrane potential and reactive oxygen species (ROS) production which are decreased by metformin. **a-d,** In representative fluorescence-activated cell sorting (FACS) plots, compared to untreated primary bone marrow-derived macrophages (BMDMs; **a**), exposure to lipopolysaccharide (LPS; **b**) or PE particles (**c**) increases mitochondrial ROS as measured by MitoSOX Red (MitoSOX); elevated mitochondrial ROS levels are decreased by metformin (met). **eh,** Similarly, mitochondrial membrane potential (measured by tetramethylrhodamine methyl ester, TMRM) of untreated BMDMs (**e**) is increased by LPS (**f**) or PE particles (**g**); increased membrane potential is decreased by metformin (**h**). **i-j**, Quantified mean fluorescence intensities (MFI) for mitoSOX (**i**) and TMRM (**j**) corroborate images in representative FACS plots. Mean (SD), n = 3, one-way ANOVA followed by Tukey’s post-hoc test.

Seeing that inhibition of complex I reduced aberrantly elevated OCR (Fig. 2c) without further decreasing bioenergetics (Fig. 1a), ECAR (Fig. 2a) or PER (Fig. 2b) in comparison to groups exposed to only polyethylene particles, we sought to determine the selective contribution of OCR from complex I to macrophage activation by polyethylene particles. Exposure to polyethylene particles increased proinflammatory cytokines, including IL-1β, IL-6, MCP-1 and TNF-a in comparison to untreated cells (Fig. 5a-d)^22^. Levels of IL-13 and IFN-g were unchanged (data not shown). Metformin decreased elevated IL-1β (Fig. 5a), IL-6 (Fig. 5b) and MCP-1 (Fig. 5c) but not TNF-a protein levels (Fig. 5d) when compared to groups exposed to only polyethylene particles. Furthermore, metformin neither increased IL-4 (Fig. 5e) nor IL-10 (Fig. 5f) protein expression in comparison to groups treated with only polyethylene particles.

**Figure 5.**
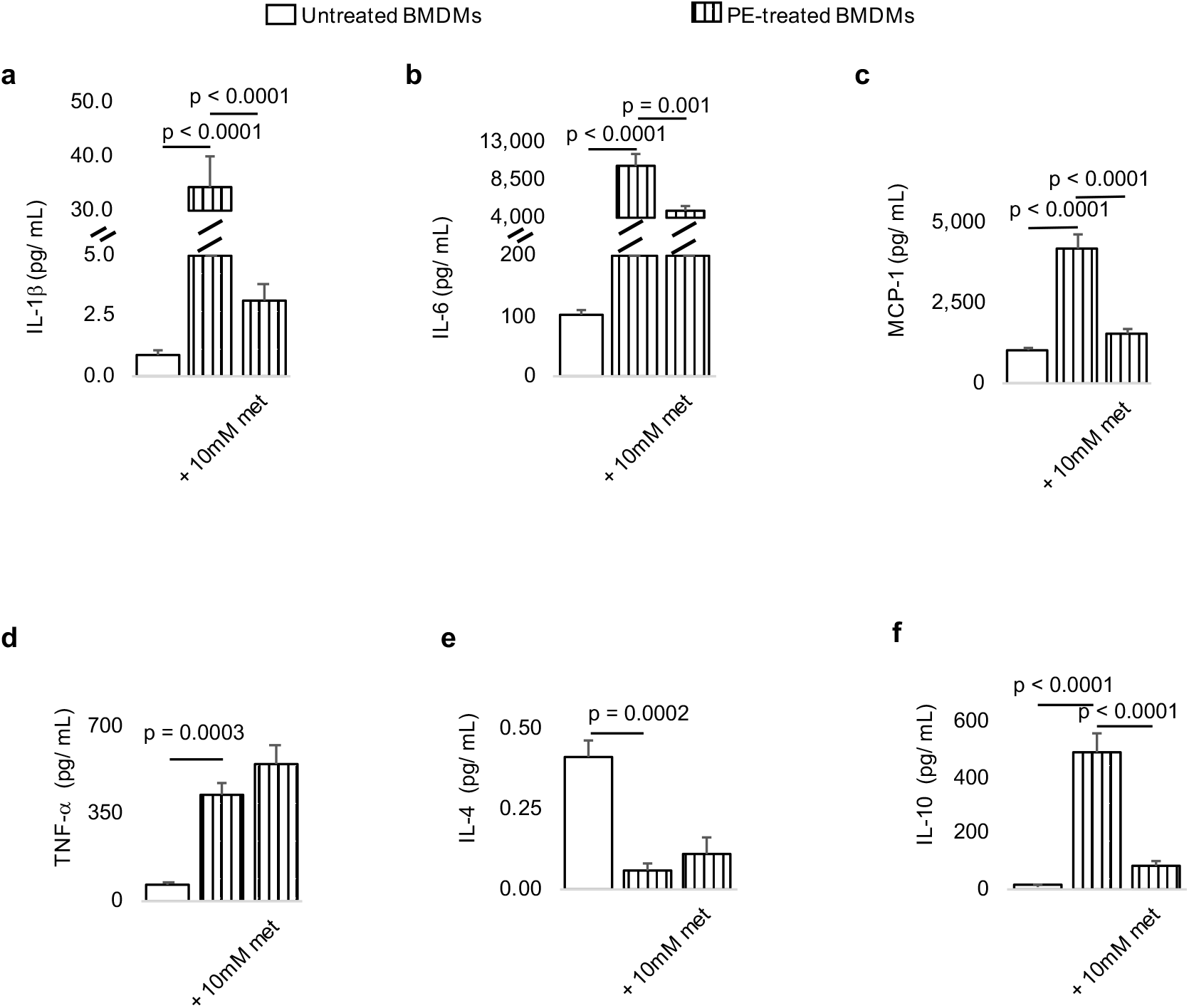
Exposure of macrophages to ultrahigh molecular weight polyethylene (PE) particles elevates proinflammatory cytokines which are decreased by metformin. **a-c,** In primary bone marrow-derived macrophages (BMDMs), proinflammatory cytokine (protein) levels, including IL-1β (**a**), IL-6 (**b**) and MCP-1 (**c**) are increased by exposure to PE particles, however, inhibition of the mitochondrial electron transport chain by metformin (met) decreases proinflammatory cytokine levels. **d**, Although the proinflammatory cytokine TNF-a is increased by exposure to PE particles, metformin does not decrease its levels. **e-f**, Whereas PE particles decrease IL-4 protein expression (**e**), they increase IL-10 levels (**f**); addition of metformin does not increase either anti-inflammatory cytokine relative to groups treated with PE particles alone. Mean (SD), n=3, one-way ANOVA followed by Tukey’s post-hoc test.

## 4. Discussion

When immune cells are exposed to non-degradable UHMWPE particles, bioenergetic (ATP) levels decrease, but mitochondrial OXPHOS is elevated. What role OXPHOS plays in immune cellular activation by polyethylene particles is not fully understood. Within the mitochondrial inner membrane and as part of OXPHOS, embedded complexes of the ETC generate ATP by electron transfer^14^. We show that OXPHOS does not contribute to ATP production in macrophages and fibroblasts exposed to polyethylene particles. Whereas inhibition of the ETC at complex I, III or V does not affect ATP production, inhibition of various glycolytic steps in immune cells exposed to polyethylene particles reduces ATP levels in a dose-dependent manner^22^. This suggests that glycolysis is primarily responsible for ATP production in immune cells exposed to polyethylene particles. On the other hand, hydrolytic degradation products of polylactide (PLA) implants aberrantly elevate ATP levels^33^. Bioenergetic alterations in immune cells exposed to PLA breakdown products compromise its biocompatibility by reprogramming cellular metabolism^33^. Accordingly, the in-vivo PLA microenvironment is characterized by enhanced fluorodeoxyglucose uptake, driving inflammation; glycolytic inhibition modulates inflammation^33^.

HMGB1/ RAGE signaling is an inflammatory pathway associated with activation after exposure to polyethylene particles^34^. As part of its inflammatory role in promoting growth of pancreatic cancers, HMGB1/ RAGE signaling directly increases ATP levels by increasing mitochondrial OXPHOS^35^. In particular, elevated oxygen consumption at complex I was observed to enhance ATP production. Inhibition of complex I activity using rotenone decreased ATP production by recombinant HMGB1 in normal fibroblasts and pancreatic tumor cells; additionally, cancer cell proliferation and migration were decreased^35^. In contrast to observations in the HMGB1 / RAGE signaling pathway, oxygen consumption is reduced in macrophages exposed to LPS^19,20^. Mitochondrial function was shown to be repurposed toward increased superoxide formation at complex I^19^, generating ROS^21^. Unlike LPS, PE particles increase oxygen consumption. Similar to macrophages exposed to polyethylene particles, concomitantly elevated glycolysis and OXPHOS for immune cellular functions is observed in neutrophils, wherein oxygen consumption is directed at activation and release of neutrophil extracellular traps (termed NETosis)^36^ and superoxide formation^37^. Similarly, CD4+ T cells obtained from humans with systemic lupus erythematosus^38^, an autoinflammatory disorder, also exhibit concomitantly elevated glycolysis and OXPHOS.

In macrophages exposed to polyethylene particles, specific pharmacologic inhibition of glycolysis was accompanied by a reduction in OXPHOS^22^. Similarly, inhibition of OXPHOS at complex III reduced both OXPHOS and glycolysis, consistent with their interdependence^39^; additionally, monocarboxylate transporter (MCT) function was reduced. Proton-linked shuttle of lactate occurs via MCTs, and these transporters are emerging targets for immunomodulation^31,32^. However, their specific role in pathologies associated with polyethylene particles requires further investigation. In contrast to complex III inhibition, inhibition of OXPHOS at complex I was not accompanied by reduction in glycolysis, allowing us to probe the selective contribution of oxygen consumption at complex I to macrophage activation by polyethylene particles. Both rotenone and metformin decreased oxygen consumption at complex I in a dose-dependent manner, consistent with their known pharmacodynamics^29^.

Our findings suggest that elevated oxygen consumption at complex I of the ETC in macrophages exposed to PE particles is directed toward mitochondrial ROS production in a manner that is dependent on mitochondrial membrane potential. During inflammation, oxygen consumption at complex I leads to superoxide formation^19^. When oxygen consumption is inhibited at complex I by rotenone, a role for reverse electron transport in superoxide formation is likely^28^ and has been shown for LPS-induced responses^19^.

Inhibition of oxygen consumption at complex I using metformin decreased only some proinflammatory cytokines, including MCP-1, IL-1β and IL-6 in primary macrophages exposed to polyethylene particles. Metformin had no effect on TNF-a protein levels which are elevated in macrophages exposed to polyethylene particles. In septic models of inflammation due to LPS, inhibition of glycolysis^24^ or OXPHOS^29^ selectively decreased IL-1 β without effects on TNF-a or IL-6 expression. In addition to the ability of metformin to inhibit oxygen consumption at complex I, it could also stimulate adenosine monophosphate (AMP)-activated kinase (AMPK). Stimulation of AMPK, a prosurvival pathway often activated during starvation, has been shown to be antiinflammatory in macrophages exposed to polyethylene particles^40^ or LPS^41^. Importantly, metformin was shown to reverse bone loss that accompanies chronic inflammation to polyethylene particles^40^.

NF-kB is the master transcriptional regulator of macrophage activation by polyethylene particles^42^. The dominant NF-kB transactivating subunit called NF-kB3 (p65, encoded by the *RelA* gene) regulates mitochondrial OXPHOS in colon carcinoma cells, and silencing *RelA* results in decreased oxygen consumption^43^. Aside from being closely associated with the mitochondrion^44^, NF-kB regulates OXPHOS by increasing cytochrome c oxidase 2^45^, a complex IV subunit, in a p53-dependent manner^46^. Consistent with this notion, p53 activation increases generation of ROS^47^. Similar to colon carcinoma, elevated oxygen consumption is an emerging feature of several types of cancers where the role of OXPHOS is multipronged. For example, elevated mitochondrial biogenesis driven by PGC-1a is associated with the invasive and metastatic capabilities of breast cancer^48^. In this role, administration of only rotenone accounts for differential oxygen consumption^48^. Pancreatic cancer stem cells exhibit a unique metabolic phenotype regulated by PGC-1α and c-MYC, and they require increased oxygen consumption for survival associated with increased ATP production^49^. Administration of oligomycin accounted for differential oxygen consumption while reducing elevated ATP levels in these cells^49^. For metastasis in pancreatic ductal adenocarcinoma, increased oxygen consumption driven by COX6B2 elevated ATP production which was abolished by oligomycin^50^. Here, ATP signaling was used by purinergic pathways required for epithelial-mesenchymal transition in metastasis^50^. In prostate, colon and breast cancers, elevated OXPHOS sustains drug resistance^51–53^.

As part of their anti-inflammatory effect, macrophages could exhibit increased mitochondrial oxygen consumption. For instance, IL-4 has been shown to increase oxygen consumption^54^. However, this increment is accounted for by administration of oligomycin which inhibits ATP synthase^54^, suggesting a need for increased levels of ATP in the anti-inflammation response. Consistent with this, inhibition of OXPHOS by nitric oxide prevents, polarization of inflammatory macrophages to an anti-inflammatory phenotype by limiting mitochondrial ATP production^20^. IL-4 induces mitochondrial biogenesis in a PGC-1 β-dependent mechanism to meet the enhanced bioenergetic needs of anti-inflammatory macrophages^55^. It has been proposed that IL-4 and IL-13 enhance OXPHOS by inhibiting mTOR^56^. In this regard, IL-4 fails to induce an anti-inflammatory phenotype when mTOR is constitutively activated in a genetic model^57^. Oxymoronically, mTOR signaling is critical for OXPHOS through PGC-1 α^58^, challenging this theory and necessitating further research to reconcile the diverse roles of OXPHOS in immune cell states.

In conclusion, we show that increased oxygen consumption does not contribute to bioenergetic (ATP) levels in activated macrophages exposed to UHMWPE particles. Rather, it is directed toward mitochondrial ROS production in a manner that is dependent on mitochondrial membrane potential, suggesting a role for reverse electron transport. Inhibition of OXPHOS in a dose-dependent manner without affecting glycolysis was accomplished by targeting complex I of the ETC using either rotenone or metformin. Consequently, this decreased expression of proinflammatory cytokines, including IL-1β, IL-6 and MCP-1 but not TNF-a in primary bone marrow-derived macrophages. These results highlight the contribution of mitochondrial respiration to activation of immune cells by polyethylene wear particles, offering new opportunities that target mitochondrial respiration to control macrophage states toward desired clinical outcomes.

## Author contributions

Conceptualization, C.V.M. and C.H.C.; Methodology, C.V.M., S.B.G., A.V.M. and C.H.C.; Investigation, C.V.M., M.M.K., O.M.B, A.T. and A.V.M.; Writing – Original Draft, C.V.M.; Writing – Review & Editing, C.V.M., M.M.K., O.M.B, A.T., A.V.M., S.B.G., and C.H.C.; Funding Acquisition, C.H.C.; Resources, S.B.G. and C.H.C.; Supervision, S.B.G. and C.H.C.

## Data availability

Data generated during this study are included in this published article.

## Acknowledgements

Euthanized C57BL/6J mice were a gift from RR Neubig (facilitated by J Leipprandt and E Lisabeth) and the Campus Animal Resources at Michigan State University (MSU). Funding for this work was provided in part by the James and Kathleen Cornelius Endowment at MSU.

## References

1 Pelt, C. E. et al. Histologic, serologic, and tribologic findings in failed metal-on-metal total hip arthroplasty: AAOS exhibit selection. JBJS 95, e163 (2013).

2 Goodman, S. B., Gallo, J., Gibon, E. & Takagi, M. Diagnosis and management of implant debris-associated inflammation. Expert review of medical devices 17, 41–56 (2020).

3 Sivananthan, S., Goodman, S. & Burke, M. in Joint Replacement Technology 373–402 (Elsevier, 2021).

4 Agarwal, R. & García, A. J. Biomaterial strategies for engineering implants for enhanced osseointegration and bone repair. Adv Drug Deliv Rev 94, 53862, doi:10.1016/j.addr.2015.03.013 (2015).

5 Ho-Shui-Ling, A. et al. Bone regeneration strategies: Engineered scaffolds, bioactive molecules and stem cells current stage and future perspectives. Biomaterials 180, 143–162 (2018).

6 Tsukamoto, M., Mori, T., Ohnishi, H., Uchida, S. & Sakai, A. Highly cross-linked polyethylene reduces osteolysis incidence and wear-related reoperation rate in cementless total hip arthroplasty compared with conventional polyethylene at a mean 12-year follow-up. The Journal of Arthroplasty 32, 3771–3776 (2017).

7 Pajarinen, J. et al. Interleukin-4 repairs wear particle induced osteolysis by modulating macrophage polarization and bone turnover. Journal of Biomedical Materials Research Part A 109, 1512–1520 (2021).

8 Goodman, S. B., Pajarinen, J., Yao, Z. & Lin, T. Inflammation and bone repair: from particle disease to tissue regeneration. Frontiers in Bioengineering and Biotechnology, 230 (2019).

9 Christman, K. L. Biomaterials for tissue repair. Science 363, 340–341 (2019).

10 Eming, S. A., Wynn, T. A. & Martin, P. Inflammation and metabolism in tissue repair and regeneration. Science 356, 1026–1030 (2017).

11 Chung, L., Maestas Jr, D. R., Housseau, F. & Elisseeff, J. H. Key players in the immune response to biomaterial scaffolds for regenerative medicine. Advanced drug delivery reviews 114, 184–192 (2017).

12 Franz, S., Rammelt, S., Scharnweber, D. & Simon, J. C. Immune responses to implants–a review of the implications for the design of immunomodulatory biomaterials. Biomaterials 32, 6692–6709 (2011).

13 Li, C. et al. Design of biodegradable, implantable devices towards clinical translation. Nature Reviews Materials 5, 61–81 (2020).

14 Nolfi-Donegan, D., Braganza, A. & Shiva, S. Mitochondrial electron transport chain: Oxidative phosphorylation, oxidant production, and methods of measurement. Redox biology 37, 101674 (2020).

15 Garaude, J. et al. Mitochondrial respiratory-chain adaptations in macrophages contribute to antibacterial host defense. Nature immunology 17, 1037–1045 (2016).

16 Olive, A. J., Kiritsy, M. & Sassetti, C. (Am Assoc Immnol, 2021).

17 Sena, L. A. et al. Mitochondria are required for antigen-specific T cell activation through reactive oxygen species signaling. Immunity 38, 225–236 (2013).

18 Di Gioia, M. et al. Endogenous oxidized phospholipids reprogram cellular metabolism and boost hyperinflammation. Nature immunology 21, 42–53 (2020).

19 Mills, E. L. et al. Succinate Dehydrogenase Supports Metabolic Repurposing of Mitochondria to Drive Inflammatory Macrophages. Cell 167, 457–470.e413, doi:10.1016/j.cell.2016.08.064 (2016).

20 Van den Bossche, J. et al. Mitochondrial dysfunction prevents repolarization of inflammatory macrophages. Cell reports 17, 684–696 (2016).

21 West, A. P. et al. TLR signalling augments macrophage bactericidal activity through mitochondrial ROS. Nature 472, 476–480 (2011).

22 Maduka, C. V. et al. Glycolytic reprogramming underlies immune cell activation by polyethylene wear particles. bioRxiv, doi:https://doi.org/10.1101/2022.10.14.512318 (2022).

23 Gonçalves, R. & Mosser, D. M. The isolation and characterization of murine macrophages. Current protocols in immunology 111, 14.11. 11–14.11. 16 (2015).

24 Tannahill, G. et al. Succinate is a danger signal that induces IL-1β via HIF-1α. Nature 496, 238–242, doi:10.1038/nature11986 (2013).

25 Ip, W. E., Hoshi, N., Shouval, D. S., Snapper, S. & Medzhitov, R. Anti-inflammatory effect of IL-10 mediated by metabolic reprogramming of macrophages. Science 356, 513–519 (2017).

26 Feoktistova, M., Geserick, P. & Leverkus, M. Crystal violet assay for determining viability of cultured cells. Cold Spring Harbor Protocols 2016, pdb. prot087379 (2016).

27 Sprague, L. et al. Dendritic cells: in vitro culture in two-and three-dimensional collagen systems and expression of collagen receptors in tumors and atherosclerotic microenvironments. Experimental cell research 323, 7–27 (2014).

28 Murphy, M. P. How mitochondria produce reactive oxygen species. Biochemical journal 417, 1–13 (2009).

29 Kelly, B., Tannahill, G. M., Murphy, M. P. & O’Neill, L. A. Metformin inhibits the production of reactive oxygen species from NADH: ubiquinone oxidoreductase to limit induction of interleukin-1β (IL-1β) and boosts interleukin-10 (IL-10) in lipopolysaccharide (LPS)-activated macrophages. Journal of biological chemistry 290, 20348–20359 (2015).

30 Pålsson-McDermott, E. M. & O’Neill, L. A. Targeting immunometabolism as an antiinflammatory strategy. Cell research 30, 300–314 (2020).

31 Samuvel, D. J., Sundararaj, K. P., Nareika, A., Lopes-Virella, M. F. & Huang, Y. Lactate boosts TLR4 signaling and NF-κB pathway-mediated gene transcription in macrophages via monocarboxylate transporters and MD-2 up-regulation. The Journal of Immunology 182, 2476–2484 (2009).

32 Tan, Z. et al. in The Journal of biological chemistry Vol. 290 46–55 (2015).

33 Maduka, C. V. et al. Polylactide Degradation Activates Immune Cells by Metabolic Reprogramming. bioRxiv, doi:https://doi.org/10.1101/2022.09.22.509105 (2022).

34 Koivu, H. et al. Autoinflammation around AES total ankle replacement implants. Foot & Ankle International 36, 1455–1462 (2015).

35 Kang, R. et al. The HMGB1/RAGE inflammatory pathway promotes pancreatic tumor growth by regulating mitochondrial bioenergetics. Oncogene 33, 567–577 (2014).

36 Sack, M. N. Mitochondrial fidelity and metabolic agility control immune cell fate and function. The Journal of clinical investigation 128, 3651–3661 (2018).

37 Guthrie, L. A., McPHAIL, L. C., Henson, P. M. & Johnston Jr, R. Priming of neutrophils for enhanced release of oxygen metabolites by bacterial lipopolysaccharide. Evidence for increased activity of the superoxide-producing enzyme. The Journal of experimental medicine 160, 1656–1671 (1984).

38 Yin, Y. et al. Normalization of CD4+ T cell metabolism reverses lupus. Science translational medicine 7, 274ra218–274ra218 (2015).

39 Van Raam, B. J. et al. Mitochondrial membrane potential in human neutrophils is maintained by complex III activity in the absence of supercomplex organisation. PloS one 3, e2013 (2008).

40 Yan, Z. et al. Metformin suppresses UHMWPE particle-induced osteolysis in the mouse calvaria by promoting polarization of macrophages to an anti-inflammatory phenotype. Molecular Medicine 24, 1–12 (2018).

41 Sag, D., Carling, D., Stout, R. D. & Suttles, J. Adenosine 5’-monophosphate-activated protein kinase promotes macrophage polarization to an anti-inflammatory functional phenotype. The Journal of Immunology 181, 8633–8641 (2008).

42 Lin, T.-H. et al. NF-κB decoy oligodeoxynucleotide enhanced osteogenesis in mesenchymal stem cells exposed to polyethylene particle. Tissue engineering Part A 21, 875–883 (2015).

43 Mauro, C. et al. NF-κB controls energy homeostasis and metabolic adaptation by upregulating mitochondrial respiration. Nature cell biology 13, 1272–1279 (2011).

44 Albensi, B. C. What is nuclear factor kappa B (NF-κB) doing in and to the mitochondrion? Frontiers in cell and developmental biology, 154 (2019).

45 Matoba, S. et al. p53 regulates mitochondrial respiration. Science 312, 1650–1653 (2006).

46 Johnson, R. F., Witzel, I.-I. & Perkins, N. D. p53-dependent regulation of mitochondrial energy production by the RelA subunit of NF-κB. Cancer research 71, 5588–5597 (2011).

47 Karawajew, L., Rhein, P., Czerwony, G. & Ludwig, W.-D. Stress-induced activation of the p53 tumor suppressor in leukemia cells and normal lymphocytes requires mitochondrial activity and reactive oxygen species. Blood 105, 4767–4775 (2005).

48 LeBleu, V. S. et al. PGC-1α mediates mitochondrial biogenesis and oxidative phosphorylation in cancer cells to promote metastasis. Nature cell biology 16, 992–1003 (2014).

49 Sancho, P. et al. MYC/PGC-1α balance determines the metabolic phenotype and plasticity of pancreatic cancer stem cells. Cell metabolism 22, 590–605 (2015).

50 Nie, K. et al. COX6B2 drives metabolic reprogramming toward oxidative phosphorylation to promote metastasis in pancreatic ductal cancer cells. Oncogenesis 9, 1–13 (2020).

51 Ippolito, L. et al. Metabolic shift toward oxidative phosphorylation in docetaxel resistant prostate cancer cells. Oncotarget 7, 61890 (2016).

52 Bacci, M. et al. miR-155 drives metabolic reprogramming of ER+ breast cancer cells following long-term estrogen deprivation and predicts clinical response to aromatase inhibitors. Cancer research 76, 1615–1626 (2016).

53 Denise, C. et al. 5-fluorouracil resistant colon cancer cells are addicted to OXPHOS to survive and enhance stem-like traits. Oncotarget 6, 41706 (2015).

54 Tan, Z. et al. Pyruvate dehydrogenase kinase 1 participates in macrophage polarization via regulating glucose metabolism. The Journal of immunology 194, 6082–6089 (2015).

55 Vats, D. et al. Oxidative metabolism and PGC-1β attenuate macrophage-mediated inflammation. Cell metabolism 4, 13–24 (2006).

56 Kelly, B. & O’neill, L. A. Metabolic reprogramming in macrophages and dendritic cells in innate immunity. Cell research 25, 771–784 (2015).

57 Byles, V. et al. The TSC-mTOR pathway regulates macrophage polarization. Nature communications 4, 1–11 (2013).

58 Cunningham, J. T. et al. mTOR controls mitochondrial oxidative function through a YY1–PGC-1α transcriptional complex. Nature 450, 736–740 (2007).

